# *Candida albicans* oropharyngeal infection is an exception to iron-based nutritional immunity

**DOI:** 10.1101/2023.01.11.523704

**Authors:** Norma V. Solis, Rohan S. Wakade, Scott G. Filler, Damian J. Krysan

## Abstract

*Candida albicans* is a commensal of the human gastrointestinal tract and one of the most causes of human fungal disease, including mucosal infections such as oropharyngeal candidiasis and disseminated infections of the bloodstream and deep organs. We directly compared the in vivo transcriptional profile of *C. albicans* during oral infection and disseminated infection of the kidney to identify niche specific features. Although the expression of a set of environmentally responsive genes were correlated in the two infection sites (Pearson R^2^, 0.6), XXX genes were differentially expressed. Virulence associated genes such as hyphae-specific transcripts were expressed similarly in the two sites. Genes expressed during growth in a poor carbon source (*ACS1* and *PCK1*) were upregulated in oral tissue relative to kidney. Most strikingly, *C. albicans* in oral tissue shows the transcriptional hallmarks of an iron-replete state while in the kidney it is in the expected iron starved state. Interestingly, *C. albicans* expresses genes associated with a low zinc environment in both niches. Consistent with these expression data, deletion of two transcription factors that activate iron uptake genes (*SEF1*, *HAP5*) have no effect on virulence in a mouse model of oral candidiasis. During microbial infection, the host sequesters iron and other metal nutrients to suppress growth of the pathogen in a process called nutritional immunity. Our results indicate that *C. albicans* is subject to iron and zinc nutritional immunity during disseminated infection but is exempted from iron nutritional immunity during oral infection.

## Importance

Nutritional immunity is a response by which infected host tissue sequesters nutrients such as iron to prevent the microbe from efficiently replicating. Microbial pathogens subjected to iron nutritional immunity express specific genes to compensate for low iron availability. By comparing the gene expression profiles of the common human fungal pathogen *Candida albicans* in two infection sites, we found that *C. albicans* infecting the kidney was iron starved and, thus, subject to iron nutritional immunity. In contrast, during oral infection, *C. albicans* is in an iron replete state and thus excepted from iron nutritional immunity. Consistent with this model, transcription factors that activate iron starvation responses are not required for *C. albicans* virulence during oral infection but are required for disseminated infection of the kidney. Thus, our work provides a striking exception to nutritional iron immunity that depends on the specific infection site of *C. albicans*.

During microbial infection, mammalian hosts limit metal micronutrient availability to reduce the fitness of the infecting organism. This strategy is referred to as nutritional immunity and is an important feature of the immune response to a variety of microbial pathogens (1). For example, the human fungal pathogen *Candida albicans* shows the transcriptional signature of iron starvation during disseminated infection of the kidney (2, 3). Furthermore, strains lacking genes required for survival in low iron environments are less virulent in mouse models of disseminated infection (4). *C. albicans* is also a commensal of the human oral cavity and gastrointestinal tract. In its commensal state within the gut, *C. albicans* appears relatively iron replete based on the expression of iron-regulated transcription factor (TF) genes such as *SEF1* and *SUF1* and their target genes (4). Thus, the commensal state of *C. albicans* is relatively iron replete while, after dissemination to the kidney, it is iron-starved.

In addition to being a commensal of the oral cavity, *C. albicans* can also invade the local submucosae to cause oropharyngeal candidiasis (OPC) in susceptible patients such as those with reduced T-cell function, living with HIV/AIDS, or undergoing oral radiation therapy (5). The oral submucosa is anatomically and physiologically distinct from target organs of disseminated candidiasis such as kidney, liver and spleen (6). To characterize the effect of different infection environments on the transcriptional profiles of *C. albicans*, we performed in vivo transcriptional profiling with a set of environmentally responsive genes during infection of either mouse kidney or tongue using standard models of disseminated (2) and oropharyngeal candidiasis (7), respectively.

By regression analysis, *C. albicans* gene expression in the two infected tissues correlates reasonably well (Fig. S1, R^2^ 0.61, Pearson r). At the gene level, however, 97 genes were differentially expressed (OPC normalized to kidney; 30 genes downregulated and 67 upregulated; ±2 fold with FDR <0.1; Benjamini-Yekutieli) as summarized in the volcano plot (Fig. 1A, See Table S1 for raw data, data normalization and analysis). *C. albicans* is a dimorphic fungus that exists primarily in the hyphal state when infecting either kidney or oral tissue (8). Virulence-associated, hyphae-specific genes such as *ECE1, ALS3*, and *HWP1* are expressed comparably in the two infection sites while *HYR1* and *SAP6* are expressed higher in oral tissue (Fig. 1B). Thus, the expression profiles indicate that *C. albicans* is primarily in the invasive, hyphal morphology in both infection sites. Interestingly, expression of the yeast phase specific gene *YWP1* (9) is undetectable in the kidney but is expressed in the oral cavity. Although the expression of *YWP1* is 30-fold lower than that observed under yeast phase growth in vitro (10), this indicates that the yeast phase may be more prevalent in OPC. The pH of the infection environment can be inferred from the relative expression of the *PHR1* (alkaline) and *PHR2* (acidic) (ref. 11); as shown in Fig. 1C, the *PHR1/PHR2* expression is skewed toward the alkaline-induced *PHR1* under both conditions with the *PHR1/PHR2* ratio higher in oral infection (*PHR1/PHR2:* kidney 8.8; OPC 20.8) indicating that the *C. albicans* experiences a neutral to alkaline environment in both infection sites. Accordingly, Rim101, the TF that drives *PHR1* expression under alkaline conditions, is required for full virulence in both infection models (12).

**Fig. 1.**
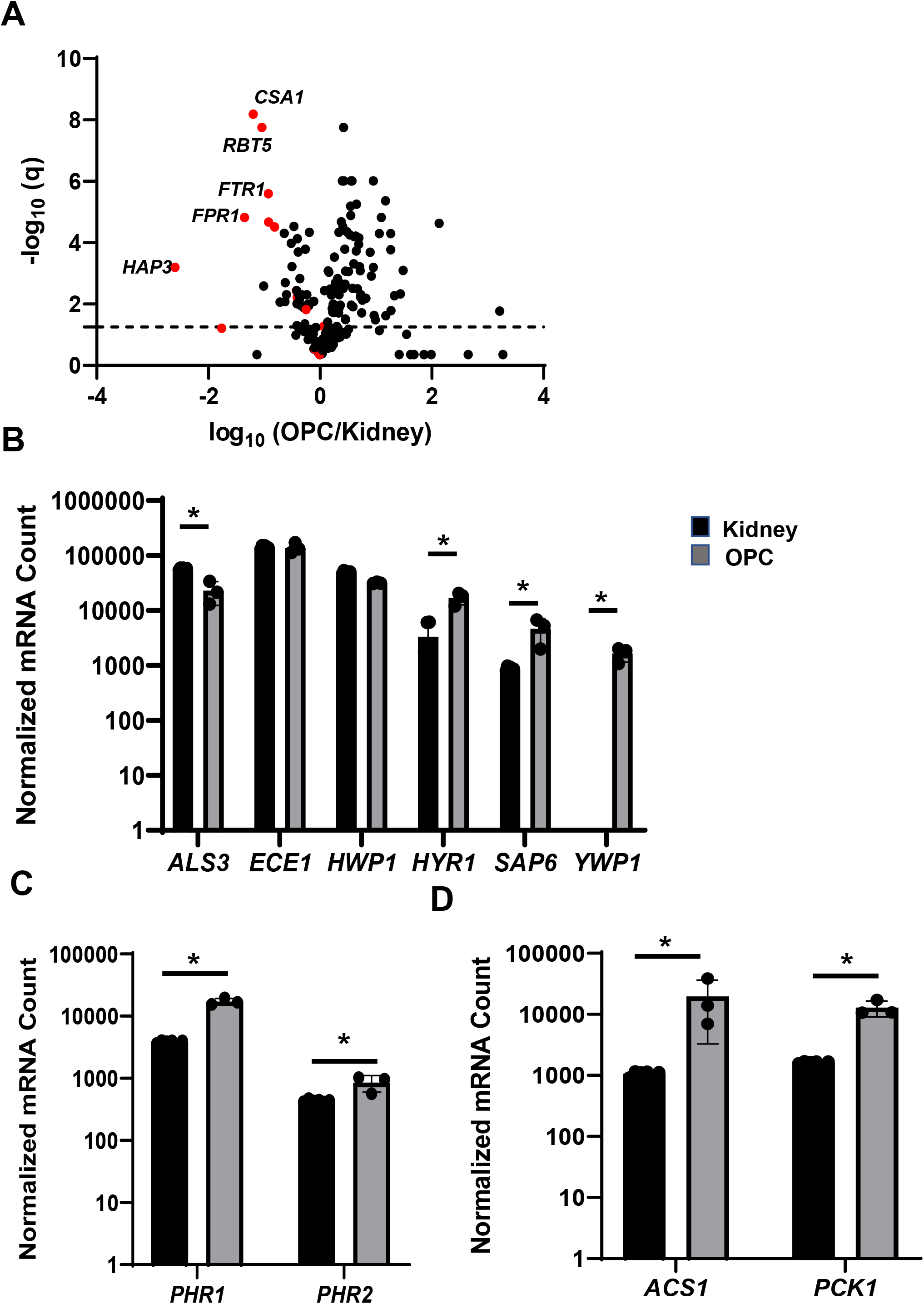
The expression of environmentally responsive and virulence genes is distinct during oral and disseminated kidney infection models. **A.** The volcano plot shows log10-fold change of genes (OPC/kidney) with the false discovery rate q (Benjamini-Yekutieli) cut-off of 0.1 shown by the horizontal line on the plot. Genes that are induced in low Fe conditions are highlighted in red. **B**. The normalized expression of hyphae-specific genes (A*LS3*, *ECE1, HWP1*, and *SAP6*) and a yeast specific gene (*YWP1*) are shown in log_10_ scale. *YWP1* was undetectable in the kidney samples. Asterisk * indicates adjusted p value (q) < 0.1. **C**. alkaline-induced gene *PHR1* is expressed at higher levels than the acid-induced *PHR2* in both oral and kidney tissue. **E**. Genes indicative of glucose starvation, *ACS1* and *PCK1*, are induced in oral tissue relative to kidney tissue.

These transcriptional data also provide insights into the relative carbon and metal nutrient status of *C. albicans* at the two infection sites. First, *PCK1* and *ACS1*, enzymes that mediate gluconeogenesis (*PCK1*) and non-glucose-derived acetyl CoA synthesis (*ACS1*), are highly expressed in oral tissue relative to the kidney (Fig. 1D). In vitro, *PCK1* (13) and *ACS1* (14) are suppressed by glucose and induced by poor carbon carbon sources such as lactate or glycerol. As such, *C. albicans* appears more dependent non-glucose carbon sources in oral tissue relative to kidney tissue.

Second, zinc is a critical micronutrient that is subject to nutritional immunity and is sequestered by the host in response to disseminated *C. albicans* infection (2), leading to expression of genes indicative of zinc starvation (Fig. 2A). *C. albicans* infecting the oral cavity expresses zinc-related genes at statistically similar levels to kidney. Third, genes induced during iron starvation (4) are expressed at dramatically lower levels (up to 30-fold) in oral tissue relative to kidney (Fig. 2B). Correspondingly, iron utilization genes that are induced in the iron replete state are also expressed at much higher levels in oral tissue. Consistent with this transcriptional signature (4, 15), TFs that regulate the expression of genes in response to iron-deficiency (*HAP2* and *HAP3*) are repressed in oral tissue while the iron-replete expressed TF *SEF2* is induced in oral tissue relative to kidney (Fig. 2C). These comparative expression data strongly support the conclusion that oral tissue is an iron-replete *C. albicans* infection site.

**Fig. 2.**
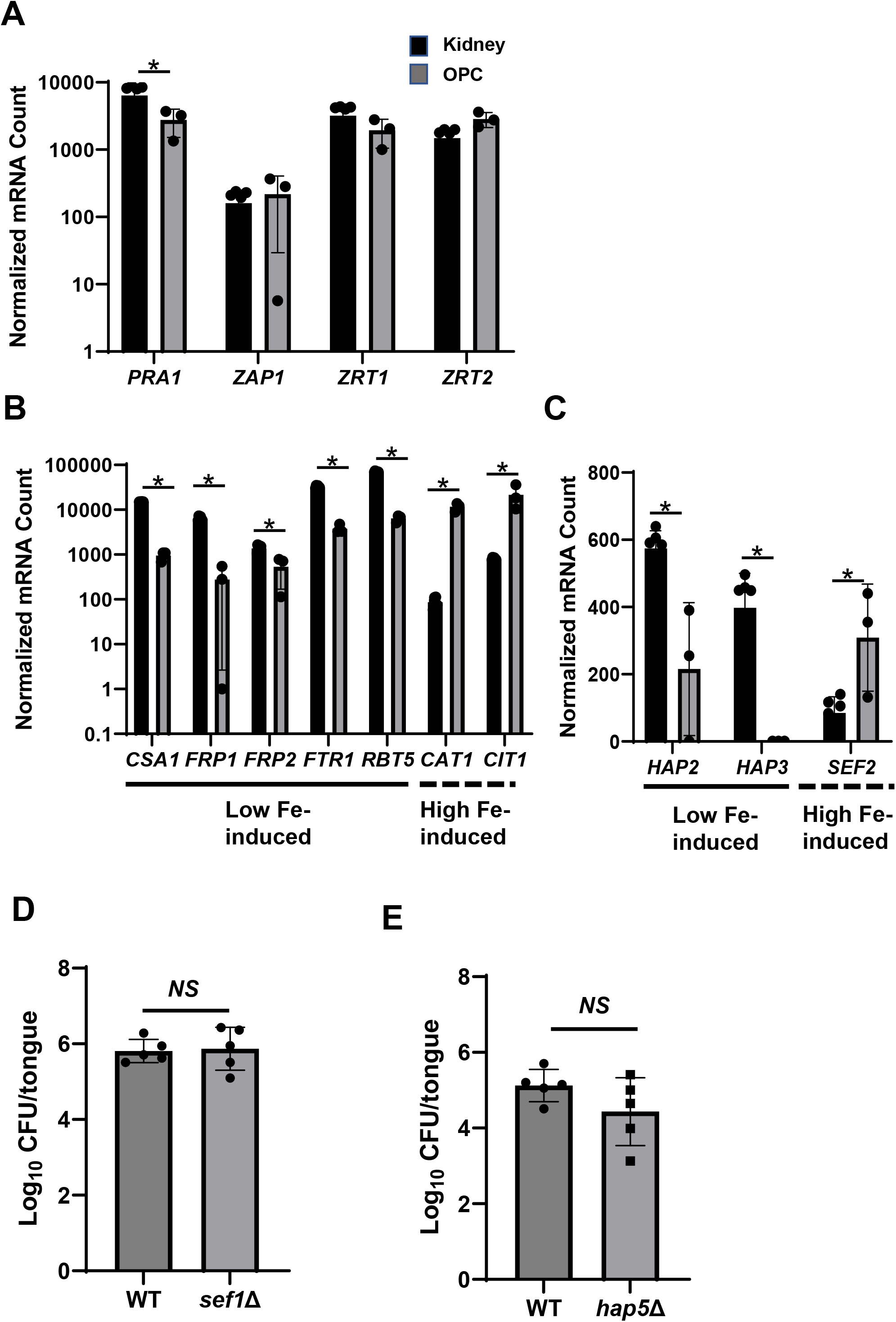
*C. albicans* does not show characteristics of iron-starvation during oral infection. **A.** The expression of zinc-responsive genes is similar in oral and kidney tissue. **B.** Genes that are induced during low iron growth (*CSA1*, *FRP1, FTR1, RBT5*) are expressed at much lower levels in oral tissue compared to kidney while genes induced in iron-replete conditions are expressed higher in oral tissue (*CAT1*, *CIT1*). Asterisk * indicates adjusted p value (q) < 0.1. **C.** Low iron-induced transcription factors *HAP2* and *HAP3* are expressed higher during infection of kidney tissue compared to oral tissue; *SEF2* is induced in iron replete conditions and is expressed higher in oral tissue. The oral fungal burden (CFU/tongue) at day 5 post infection is shown for two transcriptional regulators of the response to iron-starvation *SEF1* (**D**) and *HAP5* (**E**). The fungal burden in mice infected with the two mutants did not differ from that of mice infected with the WT reference strain (Student’s t test, P> 0.05).

If these transcriptional distinctions have pathobiological significance, then deletion of TFs required for replication in low iron environments should have no effect on OPC virulence. To test this hypothesis, we examined the virulence of strains lacking *SEF1* and *HAP5*, two TFs required for in vitro growth of *C. albicans* on low iron media. The oral fungal burden of mice infected with either the *sef1*ΔΔ or *hap5*ΔΔ mutants is not different than WT (Fig. 2D). Thus, low iron response transcriptional regulators are dispensable for *C. albicans* virulence in OPC. These genetic results are consistent with our transcriptional data and provide strong support for a model in which oral tissue is an iron replete niche for *C. albicans*.

The relative expression of environmentally responsive genes during OPC and disseminated kidney infection indicates that virulence-associated genes are expressed similarly but that the two tissues represent significantly different environments with respect to both carbon and metal nutrients. Most striking is the finding that *C. albicans* infection of the kidney appears to induce iron nutritional immunity while that of the oral cavity does not. Specific physiological features of the oral cavity and the immune response to *C. albicans* in oral tissue provide potential explanations for the lack of an iron-based nutritional immune response in this niche. First, iron is the second most abundant metal in saliva and a significant proportion of the total iron in saliva is soluble (~30%), indicating there may be a substantial pool of available iron (16, 17). Second cells of the oral mucosae lack divalent cation transporters present in other mucosal tissues and, therefore, have no known mechanism to sequester iron (18). Third, IL-17, the critical mediator of the innate immune response to *C. albicans* in oral tissue, induces the expression of lipocalin 24p3, a protein that binds to, and inhibits, catecholate-class, bacterial iron siderophores (19). Although lipocalin is induced by *C. albicans*, it does not express catecholate-siderophores and should not be susceptible to this effector of iron nutritional immunity (20). Accordingly, deletion of lipocalin in mice has no effect on OPC virulence (20). Thus, the environmental/physiological features of oral tissue and the nature of the innate immune response to *C. albicans* in that niche appear to except it from iron nutritional immunity. Taken together, these data also provide mechanistic support for the concept that treatment with exogenous iron chelators may reduce *C. albicans* replication in the oral cavity, an approach that has shown pre-clinical efficacy in mouse models (21).

Finally, this work highlights how variations in the local physiological environment of host niches impact not only pathogen physiology but also the nature and/or efficacy of the host immune response.

## Methods and materials

### General methods and strains

All *C. albicans* strains were in the SN background and have been previously reported (22). The low iron growth phenotypes for the *sef1*ΔΔ and *hap5*ΔΔ mutants were confirmed. Yeast strains were struck from frozen stocks and pre-cultured in yeast peptone dextrose medium at 30°C prior to preparation of inoculum for infection.

### Oropharyngeal candidiasis model

The immunosuppressed mouse model of OPC was employed as previously described with some modification (7). Male ICR mice were injected subcutaneously with cortisone acetate (300 mg/kg of body weight) on infection days: −1, 1, and 3. On the day of infection, the animals were sedated with ketamine and xylazine and a swab saturated with *C. albicans* strain SN250, the *sef1*ΔΔ mutant, or the *hap5*ΔΔ mutant (10^6^ cells per ml) was placed sublingually for 75 min. On post-infection day 5, the mice were sacrificed and the tongues were harvested. For fungal burden studies, the harvested tongues were homogenized and plated for quantitative fungal burden (n =5 per strain). The log10-transformed fungal burden data for each experiment was analyzed by Student’s t test to identify statistically significant differences between individual strains (*P* < 0.05). For expression studies, mice were sacrificed after 5 days of infection, and the tongues were harvested. Using a cell scraper, the *C. albicans* was scrapped off the tongue and RNA was extracted from the collected cells according to the manufacturer protocol (RiboPure RNA Purification Kit).

### Disseminated candidiasis model

As previously described (23), 5-6 weeks old, female DBA2/N mice (Envigo) were inoculated with 5 × 10^4^ CFU of SN250 by lateral tail vein injection. After 48 hrs, mice were euthanized, kidneys harvested, and placed directly into ice-cold RNA Later solution (n = 6). The kidneys were then flash frozen in liquid nitrogen and ground into a fine powder with liquid nitrogen. The resulting tissue powder is mixed with the ice-cold Trizol. The samples were placed on a rocker at room temperature (RT) for 15 min and further the cell debris were removed by centrifuged the samples at 10K rpm at 4^0^C for 10 min. Cleared Trizol was collected into a new 1.5 ml Eppendorf tube and 200 μl of RNase free chloroform was added. Tubes were shaken vigorously for 15s and kept at RT for 5 min. Further the samples were centrifuged at 12K rpm for 15 min at 4^0^C. The cleared aqueous layer then transferred to new 1.5 ml tube and RNA was further extracted following the Qiagen RNeasy kit protocol.

### Nanostring analysis

As previously reported (23), RNA (40 ng for kidney sample and 3 μg tongue sample) was added to a NanoString codeset mix (Table S1) and incubated at 65^0^C for 18 hours. After hybridization reaction, samples were proceeded to nCounter prep station and samples were scanned on an nCounter digital analyzer. NCounter .RCC files for each sample were imported into nSolver software to evaluate the quality control metrics. Using the negative control probes the background values were first assessed. The mean plus standard deviation of negative control probes value was defined and used as a background threshold and this value is subtracted from the raw counts. The background subtracted total raw RNA counts were normalized against the highest total counts from the biological triplicates. The statistical significance of changes in counts was determined by two-tailed Student’s t test (p <0.05) followed by correction for multiple comparisons using the Benjamini-Yekutieli procedure and a false discovery rate or q value of 0.1. The expression data are summarized in Table S1. Probes that were below background were set to a value of 1 to allow statistical analysis. The raw counts, normalized counts, and statistical analyses are also provided in Table S1. The data for the kidney infection model was previously reported (23).

## Acknowledgements

This work was supported by NIH grants R01AI133409 (DJK) and and R01DE026600 (SGF).

## Author Contributions

Conceptualization: Damian J. Krysan, Scott G. Filler

Formal analysis: Norma V. Solis, Rohan W. Wakade, Scott G. Filler Damian J. Krysan

Investigation: Norma V. Solis, Rohan S. Wakade

Methodology: Norma V. Solis, Rohan S. Wakade,

Supervision: Scott G. Filler, Damian J. Krysan

Writing-original draft: Damian J. Krysan

Writing-review and editing: Scott G. Filler, Damian J. Krysan

## Supplementary Figure and Table legends

**Fig. S1.**
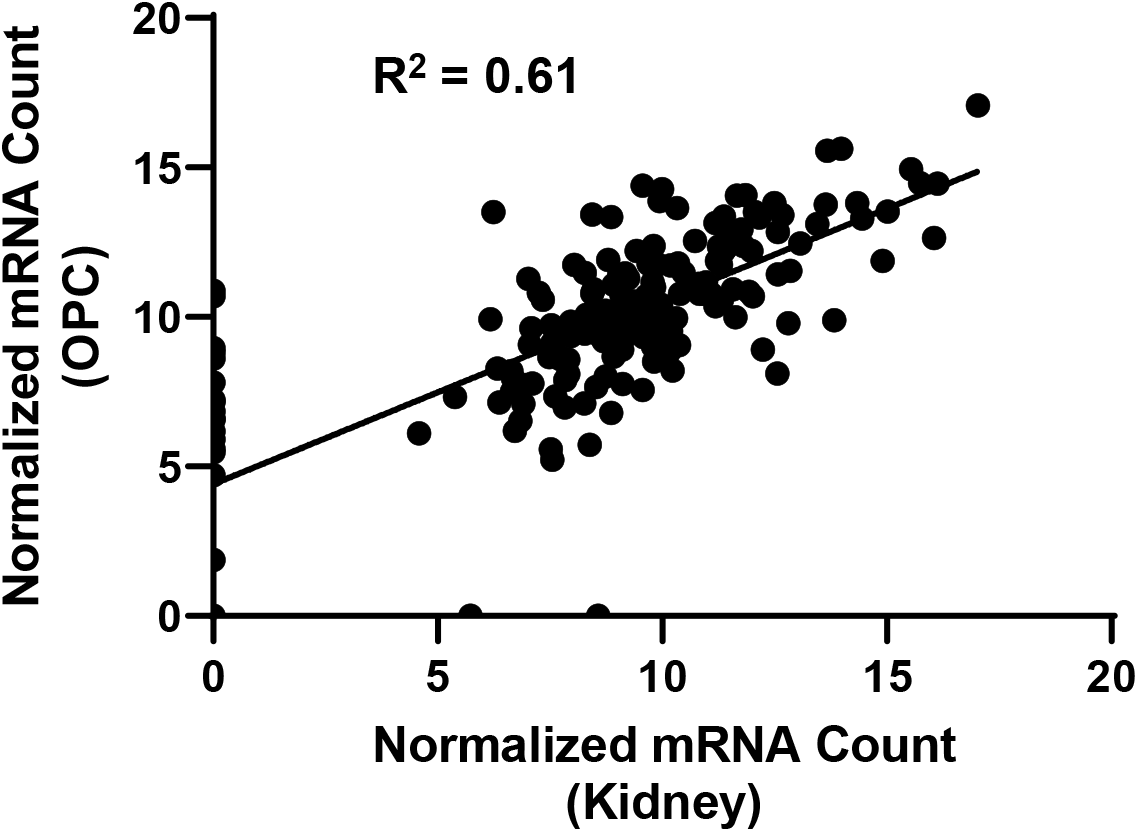
Correlation between expression of environmentally responsive *C. albicans* genes during oral and disseminated kidney infection. The normalized expression of environmentally responsive genes evaluated by nCounter correlate between *C. albicans* within oral tissue (OPC) and kidney tissue with Pearson R^2^ =0.61.

**Table S1**. Expression of environmentally responsive *C. albicans* genes in kidney during disseminated infection and tongue from oropharyngeal infection. The raw counts for kidney (six replicates) and tongue (three replicates); normalized counts; mean; standard deviation; fold change OPC relative to kidney; p value (Student t test); and adjusted p value (q) from Benjamini-Yekutieli procedure are shown. Green are genes upregulated 2-fold and red are genes downregulated 2-fold with false discovery rate (q) < 0.1.

## References

1. Murdoch CC, Skaar EP. 2022. Nutritional immunity: the battle for nutrient metals at the host-pathogen interface. Nat Rev Microbiol. 20:657–670.

2. Xu W, Solis NV, Ehrlich RL, Woolford CA, Filler SG, Mitchell AP. 2015. Activation and alliance of regulatory pathways in *C. albicans* during mammalian infection. PLoS Biol 13:e1002076.

3. Hebecker B, Vlaic S, Conrad T, Bauer M, Brunke S, Kapitan M, Linde J, Hube B, Jacobsen ID. 2016. Dual-species transcriptional profiling during systemic candidiasis reveals organ-specific host-pathogen interactions. Sci Rep. 6:36055.

4. Chen C, Pande K, French SD, Tuch BB, Noble SM. 2011. An iron homeostasis regulatory circuit with reciprocal roles in *Candida albicans* commensalism and pathogenesis. Cell Host Microbe. 10:118–135.

5. Vila T, Sultan AS, Montelongo-Jauregui D, Jabra-Rizk MA. 2020. Oral candidiasis: a disease of opportunity. J Fungi (Basel). 6:15.

6. Fanning S, Xu W, Solis N, Woolford CA, Filler SG, Mitchell AP. 2012. Divergent targets of *Candida albicans* biofilm regulator Bcr1 in vitro and in vivo. Eukaryot Cell. 11:896–904.

7. Solis NV, Filler SG. 2012. Mouse model of oropharyngeal candidiasis. Nat Protoc. 7:637–642.

8. Sudbery PE. Growth of *Candida albicans* hyphae. Nat Rev Microbiol. 2011;9(10):737–48.

9. Granger BL, Flenniken ML, Davis DA, Mitchell AP, Cutler JE. 2005. Yeast wall protein 1 of *Candida albicans*. Microbiology (Reading). 151:1631–1644.

10. Wakade RS, Kramara J, Wellington M, Krysan DJ. 2022. *Candida albicans* filamentation does not require the cAMP-PKA pathway in vivo. mBio. 13:e0085122.

11. Davis D, Wilson RB, Mitchell AP. 2000. RIM101-dependent and -independent pathways govern pH responses in *Candida albicans*. Mol Cell Biol. 20:971–978.

12. Nobile CJ, Solis N, Myers CL, Fay AJ, Deneault JS, Nantel A, Mitchell AP, Filler SG. 2008. *Candida albicans* transcription factor Rim101 mediates pathogenic interactions through cell wall functions. Cell Microbiol. 10:2180–2196.

13. Leuker CE, Sonneborn A, Delbrück S, Ernst JF. 1997. Sequence and promoter regulation of the *PCK1* gene encoding phosphoenolpyruvate carboxykinase of the fungal pathogen *Candida albicans*. Gene. 192:235–240.

14. Carman AJ, Vylkova S, Lorenz MC. 2008. Role of acetyl coenzyme A synthesis and breakdown in alternative carbon source utilization in *Candida albicans*. Eukaryot Cell. 7:1733–1741.

15. Singh RP, Prasad HK, Sinha I, Agarwal N, Natarajan K. 2011. Cap2-HAP complex is a critical transcriptional regulator that has dual but contrasting roles in regulation of iron homeostasis in *Candida albicans*. J Biol Chem. 286:25154–25170.

16. Norris HL, Friedman J, Chen Z, Puri S, Wilding G, Edgerton M. 2018. Salivary metals, age, and gender correlate with cultivable oral *Candida* carriage levels. J Oral Microbiol 10:1447216.

17. Hong JH, Kim KO. 2001. Operationally defined solubilization of copper and iron in human saliva and implications for metallic flavor perception. Eur Food Res Technol 233:973–983.

18. Sharp P, Tandy S, Yamaji S, Tennant J, Williams M, Singh Srai SK. 2002. Rapid regulation of divalent metal transporter (DMT1) protein but not mRNA expression by non-haem iron in human intestinal Caco-2 cells. FEBS Lett 510:71–76.

19. Liu Z, Peterson R, Devireddy L. 2013. Impaired neutrophil function in 24p3 null mice contributes to susceptibility to bacterial infections. J Immunol 190:4692–4706.

20. Ferreira MC, Whibley N, Mamo AJ, Siebenlist U, Chan YR, Gaffen SL. 2014. Interleukin-17-induced protein lipocalin 2 is dispensable for immunity to oral candidiasis. Infect Immun. 82:1030–1035.

21. Puri S, Kumar R, Rojas IG, Salvatori O, Edgerton M. 2019. Iron chelator deferasirox reduces *Candida albicans* invasion of oral epithelial cells and infection levels in murine oropharyngeal candidiasis. Antimicrob Agents Chemother. 63:e02152-18.

22. Homann OR, Dea J, Noble SM, Johnson AD. A phenotypic profile of the *Candida albicans* regulatory network. PLoS Genet. 5:e1000783.

23. Wakade RS, Kramara J, Wellington M, Krysan DJ. 2022. *Candida albicans* filamentation does not require the cAMP-PKA pathway in vivo. mBio. 13:e0085122.

